# Agar concentration affects the evolution of antibiotic resistance and population genomics in experimental populations *Pseudomonas aeruginosa*

**DOI:** 10.1101/2024.09.02.610736

**Authors:** Mahfuza Akter, Susan F. Bailey

## Abstract

Bacteria live in a diversity of spatially structured environments, which can impact their evolutionary dynamics via local interactions and environmental variation. Spatial structure is expected to slow the rate of adaptive evolution, increase the amount of clonal competition and so increase diversity of evolutionary trajectories explored as a population evolves. In the lab, agar is a common way in which we add spatial structure to bacterial growth environments. In this study we explored the effects of agar concertation on experimental populations of *Pseudomonas aeruginosa* evolved in the presence/ absence of a sub-lethal concentration of an antibiotic, ciprofloxacin. We varied agar across four different concentrations which modified viscosity and so the rate at which bacteria move through their environment, as well as potentially shifting the mode of motility. We saw that increasing agar concentration decreased the rate of adaptation, and that the presence of antibiotics, amplified this effect. The number and frequency of evolved mutations also varied with agar concentration and the direction of the effect changed in the presence/ absence of antibiotics – for example, the number of high frequency mutation increased with agar concentration when antibiotics were absent but decreased when antibiotics were present. We also saw an increase in the degree of parallel evolution in populations evolved in the presence of antibiotics and even more so in higher concentrations of agar. Thus, we show that agar concentration, and so spatial structure, can be an important driver of evolutionary dynamics with important impacts on antibiotic resistance evolution including the rate and predictability of adaptation.

## Introduction

Bacteria live in natural environments that are structured in many ways and across many different scales, from the detritus of a forest floor to the lungs of a human body. Even in the absence of externally imposed heterogeneities, bacteria generate their own intrinsic spatial population structure that arises from local variations in movement, competition for resources, growth rate, and other local processes. Complex bacteria population structure may even result in the formation of biofilms - communities of interacting sub-populations each with different local conditions within the population as a whole (Wimpenny et al. 2000; Lee et al. 2005), often in response to environmental stresses including limited nutrients and antimicrobial agents (Muhammad et al. 2020). Both extrinsic and intrinsically generated spatial structures are ubiquitous and so understanding how structure impacts the evolutionary dynamics of bacteria populations is important.

### Agar concentration and spatial population structure

Agar is a commonly used way to provide spatial structure for bacteria populations grown in the lab. Variation in the concentration of bacteriological agar in a growth media impacts the viscosity of the growth environment without affecting other key environmental attributes impacting population growth such as food availability, pH, or salt concentration. Depending on the concentration of agar, the particular species/ strain of bacteria, and other environmental factors (e.g. concentration of NA^+^; Terahara et al. 2017), bacteria move at a range of different speeds via a range different motility modes (e.g. swimming, swarming, twitching, sliding; Köhler et al. 2000; Overhage et al. 2008), by utilizing a variety of different appendages (e.g. flagella and pili) and secretions (e.g. exopolysaccharides) (Wadhwa & Berg 2022). Previous experiments tracking motile bacteria have identified a negative correlation between agar concentration (and so viscosity of the environment) and movement rates (e.g. *Escherichia coli:* Croze et al. 2011, *Pseudomonas aeruginosa:* Mitchell and Wimpenny 1997; Doyle et al. 2004; Kümmerli et al. 2009).

When motility rates are very low (i.e. in a high-concentration agar environment), we expect a population to effectively function as a collection of more or less independent sub-populations. Here, individuals likely only interact with other individuals in their local neighborhood. In contrast, when motility rates are very high (i.e. in a low-concentration agar or even no-agar environment) we expect a population to be unstructured or well-mixed and at the extreme, each individual will have an equal probability of interacting with all other individuals in the population. Previous experiments have shown that the result of local (as opposed to global) interactions in spatially structured populations, is often an evolved increased in phenotypic and genomic diversity (Korona et al. 1994; Rainey and Travisano 1998), and a slower rate of adaptation (Habets et al. 2006; Perfeito et al. 2008) compared to unstructured, well-mixed populations. These differences arise from spatial structure inhibiting adaptation (Kryazhimskiy et al. 2012), meaning that beneficial mutations take longer to rise to fixation, deleterious mutations take longer to be lost, and transient neutral/ nearly neutral diversity is maintained for longer.

When beneficial mutations take a long time to fix, it also becomes more likely that multiple beneficial mutations are present at the same time and so compete with each other within their local neighborhoods, known as clonal competition or clonal interference (Gerrish and Lenski 1998). In structured populations, clonal competition is expected to drive the fixation of larger effect beneficial mutations, on average, compared to mutations that fix in unstructured environments where there is less clonal competition (Martens et al. 2011). We can consider these expectations within the context of the fitness landscape concept (Kauffman and Levin 1987). If the fitness landscape is rugged (which may be typical with bacteria; e.g. Papkou et al. 2023), populations with multiple low frequency mutations present simultaneously (i.e. clonal competition in structured populations) will have the potential to explore many evolutionary trajectories at once. A multi-trajectory exploration of the fitness landscape increases the chances of a population gaining access to the highest fitness regions in a rugged landscape. In the long run, this multi-trajectory exploration may lead to an increased extent of adaptation (Nahum et al. 2015) and more similar evolutionary outcomes (parallel evolution) compared to populations evolving in unstructured, well-mixed environments with less clonal competition (Gordo and Campos 2006).

### Other impacts of agar concentration

Thus, varying the viscosity of an experimental environment by adjusting agar concentration is likely to impact the growth and evolution of a bacteria population via changes to its *degree of spatial population structure*. However, as mentioned previously, along with *rate* of motility, bacteria typically also adjust their *mode* of motility depending on the viscosity of their environment. *Pseudomonas aeruginosa* (PA), in particular, demonstrate three major types of movements: (i) swimming mediated by flagella in a liquid environment and in low agar concentrations (typically <0.3% [wt/vol]), (ii) swarming on semi-solid (viscous) surfaces (typically 0.5 to 0.7% [wt/vol]), and (iii) twitching facilitated by type IV pili on solid surfaces (1% [wt/vol] agar) (Köhler et al. 2000; Yeung et al. 2012). Importantly, shifts in these motility modes along with other potential physiological changes in response to the viscosity of the environment also have the potential to impact on how PA and other bacteria species grow and evolve in environments of varying agar concentration, beyond simple shifts in movement rates. For example, changes in movement mode and bacteria population density have both been shown to impact sensitivity of a population to antibiotics (e.g. Lai et al. 2009), with potentially important impacts on population growth and evolution.

### Previous evolution experiments in structured environments

Previous bacterial experimental studies have used a range of approaches to explore how spatial structure can impact evolutionary dynamics. For example, multi-patch metapopulations with experimenter-imposed transfer between well-mixed patches (Kryazhimskiy et al. 2012; Nahum et al. 2015), static liquid conditions that allow for biofilm formation (Ahmed et al. 2018; Santos-Lopez et al. 2019), evolving bacteria in a synthetic lung sputum media with mucin (Wong et al. 2012), and generating spatial structure in the form of semi-solid agar (Baym et al. 2016). However, there have been no studies that test the evolutionary impacts of different degrees of spatial structure adjust via agar concentration, and test how those impacts at the phenotypic and genomic level is modified by the presence versus absence of antibiotics.

### Evolution of antibiotic resistance

Variation in spatial structure has the potential to impact the evolutionary dynamics of bacteria in response to antibiotics. Bacteria vary in their intrinsic level of resistance (Cox and Wright 2013), and in some bacterial populations, antibiotic resistance may be induced by specific growth states or environmental conditions (Fernández et al. 2011), including viscosity (e.g. Kostenko et al. 2007). There have been many efforts to better understand the circumstances under which antibiotic resistance is more or less likely to evolve in bacteria. Many studies have taken the approach of collecting and characterizing antibiotic resistant bacteria from clinical patients and the environment, and then identifying mutations in key genes that drive antibiotic resistance across a range of bacteria species (e.g. Frieri et al. 2017; Qiao et al. 2018; Larsson and Flach 2022). These studies reveal some general patterns, for example, the types of genes that tend to play a role (e.g. efflux pumps, Wright 2011; Blair et al. 2015), and the tendency for associated fitness costs (Andersson et al. 2007; Schulz zur Wiesch et al. 2010; Hernando-Amado et al. 2017). Another approach is to use experimental evolution. These types of studies identify genes and mutations that can confer resistance, and also tend to characterize details of the dynamics of bacterial populations as they evolve antibiotic resistance in different types of environments (e.g. Wong et al. 2012; Jørgensen et al. 2013). However, these studies often find discrepancies between the specific resistance genes and mutations most commonly identified, likely in part, due to differences in the characteristics of the simplified experimental environment versus clinical and other “natural” environments in which bacteria typically reside. An important way in which bacterial lab culture environments often differ from those outside of the lab, is that lab environments are less complex, and often well-mixed and homogeneous.

### Experimental evolution approaches

In this study we contrasted the evolutionary impacts of a range of agar concentrations on populations of PA grown in the presence and absence of an antibiotic (ciprofloxacin). PA is an opportunistic human pathogen, causing acute infections, as well as chronic lung infections in cystic fibrosis patients (Driscoll et al. 2007). However, even when PA infections are treated with an initially effective antibiotics (often fluoroquinolones, such as ciprofloxacin used here), rapid evolution of resistance is frequently observed (Driscoll et al. 2007; Breidenstein et al. 2011; Rehman et al. 2019). Differences in strength of selection (e.g. presence/ absence of antibiotics) are expected to drive general differences in evolutionary dynamics, particularly in the rate of adaptation, and those impacts might interact with spatial structure. Beyond these general interactions, there is also the potential for antibiotic-specific interactions with spatial structure. In *Pseudomonas aeruginosa* some of the many genes identified as being involved in determining motility mode (e,g, swimming, swarming, Köhler et al. 2000; Overhage et al. 2008)) have also been shown to impact antibiotic resistance (e.g. *orfN*) (Wong et al. 2012; Jorth et al. 2017; Laborda et al. 2022). These observed pleiotropic effects suggest that agar concentration or the viscosity environment may be particularly important in the evolutionary dynamics of PA in the presence of antibiotics.

We aimed in this study to experimentally test the impact of agar concentration and antibiotics on the evolutionary dynamics of populations of *Pseudomonas aeruginosa* (PA). We created four differently structured environments, ranging from a liquid environment (0 % agar), across increasing viscosities of semi-solid agar (0.2 %, 0.4 %, and 0.6 %). Populations growing in these environments were continuously shaken at 200 rpm, so that the liquid environment was well-mixed and thus represented as close as possible to a totally *unstructured* environment.

In this study, we saw that agar concentration impacts of the evolution of *Pseudomonas aeruginosa* (PA) populations in a number of different ways depending on the presence/absence of antibiotics. We tested replicate population grown in four environments with four different viscosities, ranging from a liquid environment (0 % agar), across increasing concentrations of semi-solid agar (0.2 %, 0.4 %, and 0.6 %). All populations were continuously shaken at 200 rpm, so that the liquid environment was truly well-mixed and thus represented as close as possible to a totally *unstructured* environment. We saw that evolved changes in antibiotic resistance did not vary in a consistent way across agar concentration, however rate of adaptation and the number of evolved mutations did. Adaptation decreased with increasing agar concentration and this pattern was stronger in the presence of antibiotics. The number of putatively beneficial mutations decreased with increasing agar concentration in populations evolved in the absence of antibiotic, while in the presence of antibiotic, the number of putative beneficial mutations increased with agar concentration. The presence of antibiotics also drove the evolution of decreased motility. Our results suggest that variation in agar concertation, and so spatial structure, can have important impacts on evolutionary dynamics at the phenotypic and genomic level, and both the strength and direction of these impacts can change in the presence strong selection from an antibiotic.

## Methods

### Bacterial strains, media, and reagents

We used clonal isolates of *P. aeruginosa* PA14: LacZ+ and *P. aeruginosa* PA14:LacZ- to found 48 independent lines. These strains are isogenic to each other with the insertion of the lacZ gene (PA14: LacZ), a selectively neutral marker (as in (Wong et al. 2012). Colonies with lacZ are blue on agar plates supplemented with 40 mg/l of 5-bromo-4-chloro3-indolylbeta-D-galactopyranoside (X-Gal) and can easily be distinguished from the pale-yellow colonies of the unmarked strain. All strains and evolving populations were frozen at -80 °C in 15 % (v/v) glycerol. Unless mentioned otherwise, all initial liquid culturing was performed in King’s Broth (KB) maintained at 200 RPM of continuous shaking, and agar plate colony growth was performed on Luria-Bertani Agar (LBA), always at 37°C. Selection experiment was performed in M9 minimal salts (1 g/l NH4Cl, 3 g/l KH2PO4, 0.5 g/l NaCl, and 6.8 g/l Na2HPO4 supplemented with 15 mg/l CaCl2, 0.5 g/l MgSO4) and a carbon source, xylose (2 M) (termed as minimal media). Spatial structure were maintained using four different agar concentrations (0 %, 0.2 %, 0.4 %, and 0.6 %) representing a range of viscosities similar to those PA may experience outside of the lab (e.g. 0.5% agar viscosity: ∼ 1 mPa.s, (Yu et al. 2020), compared lung mucus viscosity ∼ 1 mPa.s, (Abrami et al. 2024). Antibiotic ciprofloxacin (Cip) stock prepared with water was stored in 4°C and all culture dilutions were performed using 0.9 % saline.

### Quantifying ancestor sensitivity to antibiotic in semi-solid agar

The sensitivity of the ancestor PA (both LacZ+ and LacZ-) to the antibiotic ciprofloxacin (Cip) was determined following a modified version of the standard minimum inhibitory concentration (MIC) method described by the Clinical and Laboratory Institute (Ma 2006). Briefly, the wells in the last column of a flat-bottomed polystyrene 96-well micro-titer plate were filled with 200 µl of minimal media supplemented with one of the experimental four agar concentrations and Cip at double the greatest desired test concentration (8 µg/ml) for the assay. Serial two-fold dilutions were performed across the plate in fresh media finishing with a final volume of 200 µl containing desired Cip concentrations in each well. Overnight cultures of PA grown in KB were adjusted by dilution to 1 × 10^8^ CFU/ml (CFU: colony forming units; absorbance at 600 nm was measured and converted to CFU using a calibration curve). After adjusting the CFU/ml, 5 µl of each bacterial suspension was added to each well in a row on the MIC plate. Two columns of the micro-titer plate were used as controls: one column with growth medium only but no antibiotic was inoculated with bacterial culture to confirm normal growth of the bacteria, and another column contained only uninoculated medium with no antibiotic to detect any contamination during the experiment. To estimate accurate CFU/ml, each inoculum was serially diluted (1:10) in saline (0.9%) and quadruplicate aliquots of 10 µl were plated on LBA. Agar plates were incubated at 37°C until colonies appeared. Micro-titer plates were also incubated at 37 °C for 24 h (200 rpm) and then examined for turbidity which indicated bacterial growth. However, growth of PA in the nutrient-limited media was not enough to visualize turbidity with the naked eye, particularly in the presence of agar which added additional turbidity making it even more difficult to identify bacterial growth. Thus, each well with different Cip concentrations were serially diluted prior, plated on LBA, and incubated at 37°C until colonies appeared. In this modified MIC assay, we estimated the growth rate (*r*) of the ancestor in all selection experiment media across a range of antibiotic concentrations (0 µg/ml to 4 µg/ml) by plating and counting initial CFU (CFU*_I_*) and final CFU (CFU*_F_*), using the following equation:

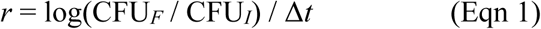

where Δ*t* is the change in time between the initial and final CFU measurements. Finally, a measure for *relative fitness* in each antibiotic concentration was estimated as the ratio of growth rate in the presence of Cip relative to growth rate in the absence of Cip. For the evolution experiment we selected the concentration of Cip resulting in approximately 50% reduction in fitness in each environment from the multiple time experiments.

### Evolution experiment

PA14:LacZ+ and PA14:LacZ- were used to found 24 independent lines each, for a total of 48 replicate populations. Replicate populations were initiated from overnight culture grown up from a single cell and transferred to two 24-well plates, with 2 ml of minimal media in each well, in an orbital shaker (200 rpm) at 37°C (∼10^8 CFU/ml). The experiment was comprised of two factors in a fully crossed design. Within one factor there were two treatment levels: 1) Cip present, and 2) Cip absent, and within the other factor there were four treatment levels comprising four different concentrations of agar – 0 % (unstructured environment), 0.2 %, 0.4 %, 0.6 % (structured environments), for a total of eight treatments. For antibiotic treated populations, we used 0.06 µg/ml Cip in the unstructured environment and 0.1 µg/ml in the structured environments to maintain a 50 % reduction in fitness in both cases (Supplementary information, Fig. S1). Each treatment contained six replicate populations, with three of those replicates inoculated with LacZ+ and other three with LacZ-, for a total of 48 replicate populations. Every 24 h (∼6.6 generations), 20 µl of each population was transferred to fresh media. This procedure resulted in population sizes fluctuating from around 10^4 up to 10^6 in the antibiotic present populations, and around 10^6 to 10^8 in antibiotic absent populations, and so effective population sizes of about 2 × 10^4 and 2 × 10^6 for the antibiotic present and absent populations, respectively (calculated as the harmonic mean). This transfer regime was repeated for 7 transfers (1 week), resulting in approximately 50 generations of selection. All evolving populations were preserved daily at -80°C in 15 % (v/v) glycerol.

Note that we chose to use a xylose-limited minimal media (a non-preferred carbon source) as the base media in this experiment. This was done in an attempt to exert significant selection pressure on the evolving populations (even in the absence of antibiotic) to allow for observation of adaptive evolution within just one week (Bailey and Kassen 2012).

### Quantifying fitness in the evolution environments

Fitness of each evolved population was estimated using growth rate in their evolution environments relative to that of the ancestor. Here, we use 24-hour growth rate as a proxy for fitness. To perform the assay, evolved populations along with the ancestors were grown up from frozen in KB media and used to inoculate 24-well plates containing minimal media without Cip. Similar to the modified MIC assay, populations were tested in the agar concentration in which they had evolved. Inoculation cultures were adjusted by dilution to 1 × 10^8^ CFU/ml before inoculation into experimental plates and cultures were plated in quadruplicate as described earlier to ensure accurate CFU counts. Experimental plates with ancestor and evolved populations were incubated at 37° C and 200 RPM shaking for 24 hours. Each well from experimental plates were then diluted and plated in LB agar plates to get the final CFU count. Growth rate over 24 hours was estimated using Eqn 1 and then relative fitness of the evolved populations was estimated as the ratio of the growth rate of the evolved population to the growth rate of the ancestor.

### Quantifying evolved antibiotic resistance

For each evolved population, level of resistance was measured as the minimal inhibitory concentration (MIC) of ciprofloxacin following standard procedure in Muller Hinton Broth (MHB) (Clinical and Laboratory Standards Institute 2009). We made one adjustment to the standard protocol, using a culture grown from a sample of the preserved evolved populations, instead of a single colony. Thus, each test culture likely contained multiple genotypes depending on the evolutionary dynamics of the population. Using this standard procedure, we observed no change in MIC for the populations evolved without antibiotic compared to the ancestor (see Fig. S2), thus we focused follow-up MIC tests on just those populations evolved *with antibiotic*. For these follow-up tests we used the modified MIC assay described above, except the Cip concentrations tested were increased, using concentrations ranging from 0.06 µg/ml to 16 µg/ml.

### Whole-genome sequencing and analysis

Genomic DNA from all 48 populations evolved for approximately 50 generations, along with the two ancestor PA strains, were extracted using a DNA extraction kit following the manufacturer’s instruction (DNeasy® Blood & Tissue Handbook). Genomic DNA was extracted from populations, as opposed to individual colonies. Whole genome sequencing of the extracted DNA was performed by Microbial Genome Sequence Center. The sequence data were quality filtered using default setting of fastp v0.20.1. After quality filtering, 96.8% of the reads were retained. SNPs and indels in the population genomes were then called using breseq v0.38.0 (Barrick et al. 2009) to predict polymorphic mutations with bowtie2 v2.5.1 and Pa14 reference genome NC_008463.1. To be called as a polymorphic variant via the breseq analysis using default parameters setting, read variations must meet a number of criteria, including being present on at a frequency of at least 5% of the reads, being present on at least two reads per strand (+/-), and statistical support via a likelihood-ratio test for the model consisting of a mixture of two alleles, versus only the reference base. Mutations that appeared in the ancestor PA compared to reference genome were removed from the evolved populations considering those mutations were already present in the ancestor, thus, unique mutations from each population were identified. After processing, we manually reviewed the resulting mutation output and noticed a number of cases where multiple neighboring INDELs had been identified as independent mutation events. We reprocessed this data to classify these cases as single multi-nucleotide INDELs. We also identified regions of the genome with high concentrations of low frequency SNPs that appeared to be associated with consistently low coverage in those regions, across independent populations. For this reason, we re-filtered the mutation data set, applying more stringent criteria (in addition to the afore mentioned breseq default criteria) for identification of variants: coverage must be greater than 75 or the variant must be present in at least 5 reads. All data presented in the results section represent the refiltered dataset, however we note that this refiltering procedure did not qualitatively change the results.

We then grouped identified evolved mutations in two ways: 1) synonymous versus non-synonymous mutations, and 2) frequency in their population: *high frequency* mutations (≥ 10 % frequency) versus *low frequency* mutations (< 10 % frequency). Our choice of a frequency cut-off of 10% is not a quantitative division between mutations driven by selection versus genetic drift, however we do suggest that the “high frequency” mutations have more clearly escaped genetic drift and so are more likely to have beneficial fitness effects compared to the “low frequency” mutations. The “low frequency” mutations are certainly still filtered by selection to some extent, but we suggest they are more likely to range in their fitness effects from beneficial to neutral, to perhaps even deleterious. Further support for examining the “high”/ “low” frequency classes separately comes from visual examination of the frequency distributions of mutations in our populations, where we see what loosely appears to be two frequency groups of mutations –indicated by one peak at very low frequencies and then another at higher frequencies with a very long tail reaching to all the way to 1 (see Fig S3).

### Estimating parallel evolution

We quantified parallel evolution or the similarity in genetic changes at the gene-level between pairs of population using the Jaccard Index, J (as in Wong et al. 2012; Bailey et al. 2015). Given two sets, G1 and G2, that list mutation-bearing genes found in populations 1 and 2, respectively,

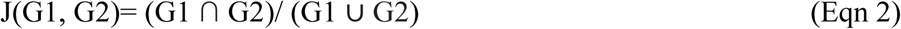

That is, J is the number of genes mutated in both populations divided by the total number of unique genes mutated in population 1 or 2. J ranges from 0 to 1, with 1 indicating identical genotypes and 0 indicating no shared mutations. We calculated J for all pairs of evolved populations. We calculated null expectations for J by using a Poisson process (‘rpois’ in R) to generate sets of random mutations across a genome (at the same rate as in the observed data), and calculating J-values for pairs of those random mutation sets. Observed J-values were then compared with null expectations to determine whether the experimental populations showed more parallel evolution than expected by chance alone.

Due to the nonindependence of J estimates, statistical significance of differences in J values from populations pairs in different treatments (e.g. antibiotic absent/ present), were estimated using a resampling procedure to generate a null distribution specific to each pairwise comparison. For each population, we resampled mutations from the set of all observed mutations and the number of mutations was set equal to the number of actual observed mutations in that population. J was recalculated for each comparison and the procedure was repeated 1000 times for each comparison, generating distributions of J values. Using these distributions, we calculated the probability of obtaining the observed J values, or greater, by chance alone (P value). Significance was assigned to probabilities less than 0.0033 – an experiment-wide significance threshold of 0.05, Bonferroni-adjusted for all comparison sets.

### Statistical analyses

All the statistical analyses of data from phenotypic and genomic assays were performed using R (version 4.3.2). Unless otherwise noted, the data were first checked for normality. If the data were not normally distributed, we performed a log transformation and tested again for normality. Data that fit the assumption of normality were analyzed using the appropriate parametric pairwise t-tests and ANOVAs. However, if the data were still not normally distributed, we used permutation ANOVA tests with the N = 1000 permutations. If the P-value estimated from the N = 1000 permutations was zero, we report it as P < 0.001 (i.e. 1/N).

## Results

### Fitness and phenotype evolution

All evolved populations had increased fitness relative to their ancestor in their evolution environments (Fig. 1). In presence of antibiotics, populations reached a higher relative fitness, while in the absence of antibiotic, populations still adapted significantly but to a lesser extent. Thus, we observed a significant effect of presence/ absence of antibiotic on fitness of evolved populations in their evolution environment (ANOVA, main effect of antibiotic, P <0.0001; see Table S1). The main effect of agar concentration was also significant (ANOVA, main effect of structure, P = 0.009; see Table S1), with relative fitness decreasing as agar concentration increased. In the presence of antibiotic, relative fitness decreased quite substantially with increasing agar concentration in the environment, while populations evolved in absence of antibiotic showed a slower decrease (ANOVA, agar conc. x antibiotic, P = 0. 006; see Table S1). A result of this interaction is that in the higher agar concentration environments (0.4% and 0.6 % agar) the presence/ absence of antibiotics does not significantly affect fitness, while in the lower concentration environments it does. We did not observe a significant relationship between change in resistance and fitness without the antibiotic (Fig. S4), and so no evidence for an evolved cost of resistance.

**Figure 1:**
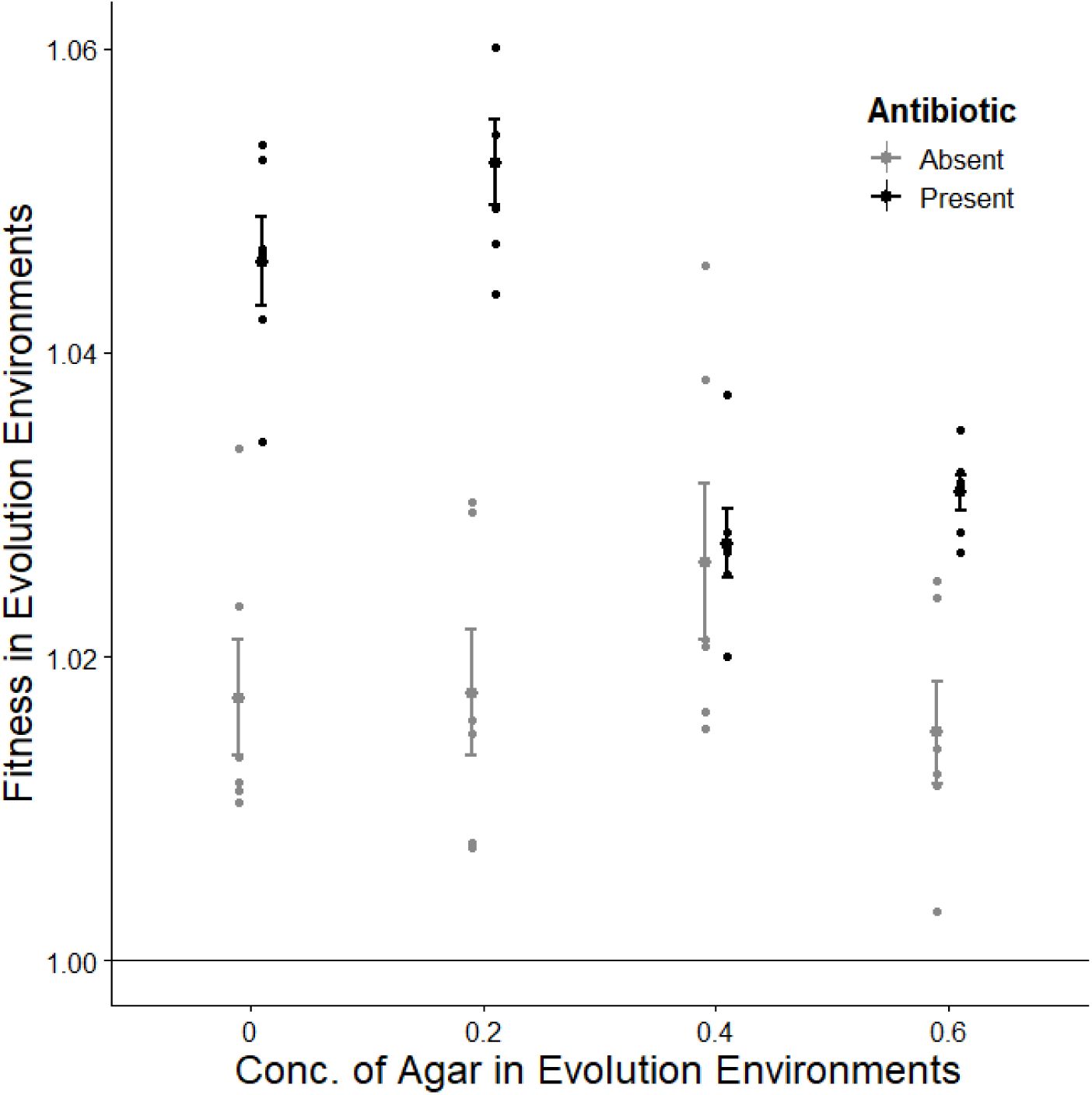
Relative fitness of populations evolved with antibiotic absent (gray) and present (black) across different concentrations of agar in their evolution environments. Scattered points represent relative fitness of each replicate, big points with error bars represent the mean of six replicate populations +/- SE. The solid horizontal line at y = 1 indicates the fitness of the ancestor.

Our modified MIC assay showed that all populations evolved in the presence of antibiotics had increased resistance to ciprofloxacin (Cip) in their evolution environment conditions (Fig. 2). In general, populations evolved in presence of Cip showed approximately 10- fold to 45-fold increase in resistance over their ancestor. The largest fold increase in resistance was observed in populations evolved in the unstructured environment (0 % agar), while smallest increase was observed in populations evolved in the highly-structured environment (0.6 % agar). However, analyzed together there was no significant effect of agar concentration on resistance (Kruskal-Wallis t-test, agar conc. P-value = 0.1106).

**Figure 2:**
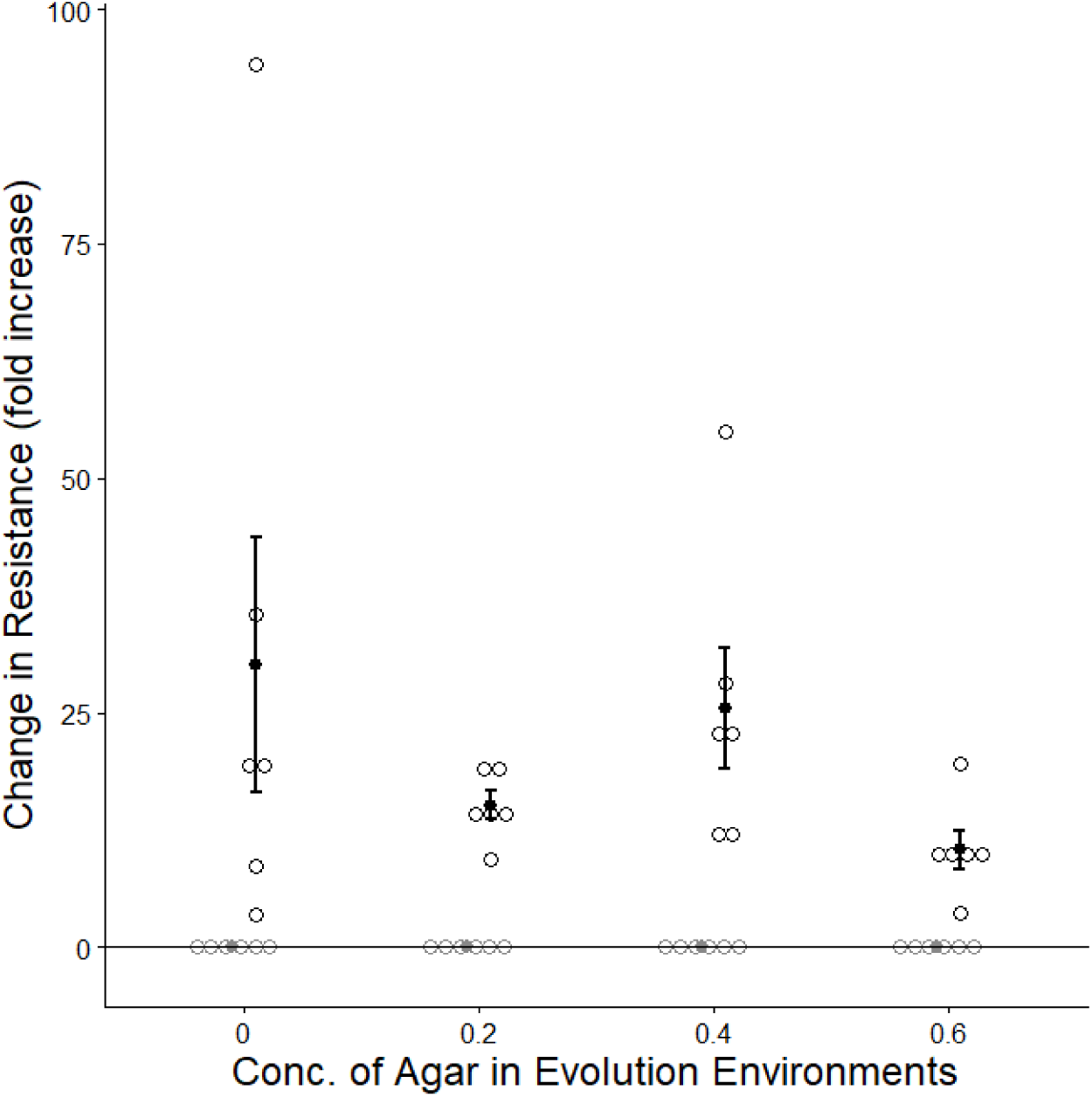
Change in resistance to ciprofloxacin of evolved populations relative to ancestor in same concentrations of agar (%) and media as their evolution environment. Scattered blank circles represent the change of resistance of each replicate, dark points with error bars represent the mean of six replicate populations +/- SE.

### Genomic evolution

Mean coverage in genome sequence data from across all 48 replicate populations and the ancestor was 147.5 (range 75.8 – 202) with a mean quality score of ∼ 98 %. Across all 48 evolved populations, the per population mean (+/- SE) number of observed mutations (at all frequencies) was 15.9 +/- 0.6 nonsynonymous, 3.5 +/- 0.3 synonymous, and 10.3 +/- 0.6 intergenic. Mean mutation numbers, grouped by treatment and mutation type are summarized in Table S2. Most of the observed mutations had not fixed. Out of the total 48 evolved populations, 19 had at least one fixed mutation (39.6%), of which only 10 populations had a fixed nonsynonymous mutation (21%). Mean number of fixed mutations per population, grouped by treatment and mutation type is summarized in Table S2. The majority of genes baring mutations were hit just once across all 48 evolved populations (∼85 %), while about 15 % of the genes baring mutations were hit in more than one independently evolved population (i.e. parallel evolution). Figure S5 summarizes the distribution of number of observed mutations per gene, across all mutations, nonsynonymous mutations alone, and synonymous mutations alone.

Across the 48 evolved populations, we identified mutations in six genes identified as antibiotic resistance genes in the NCBI antimicrobial resistance (AMR) reference gene catalogue. Those genes are: amrR, mexR, mexS, nalC, nalD, and nfxB. Mutations in these six genes were seen in 22 out of the 24 antibiotic-present populations, and in none of the antibiotic- absent populations. The majority of the antibiotic-present populations had a mutation in just one of these known AMR gene (15 out of 24), but some antibiotic-present populations had mutations in two or three different AMR genes (5 and 1 populations, respectively). The most common AMR gene to mutate across all evolved populations in this experiment was nfxB, observed in 15 of the 24 antibiotic-evolved populations.

The number of evolved mutations varied depending on both the presence/ absence of antibiotic and agar concentration (Fig. 3). For the high frequency mutations (> 10 %), their number *increased* with increasing agar concentration in populations evolved in the presence of antibiotic, however in absence of antibiotic, the number of high frequency mutations *decreased* with increasing agar concentration (Fig. 3A). A significant interaction between antibiotic and agar concentration confirms this observation (ANOVA, antibiotic x agar conc., P = 0.0009; see Table S3). In the unstructured (0 %), low (0.2 %), and medium (0.4 %) agar concentration environments, there is no significant difference in the total number of high frequency mutations in populations evolved in the presence/ absence of antibiotic. However, for high agar concentration (0.6 %) environments there *is* a significant difference between populations evolved in presence/ absence of antibiotic (P = 0.00147, Holm-adjusted pairwise T-test).

**Figure 3:**
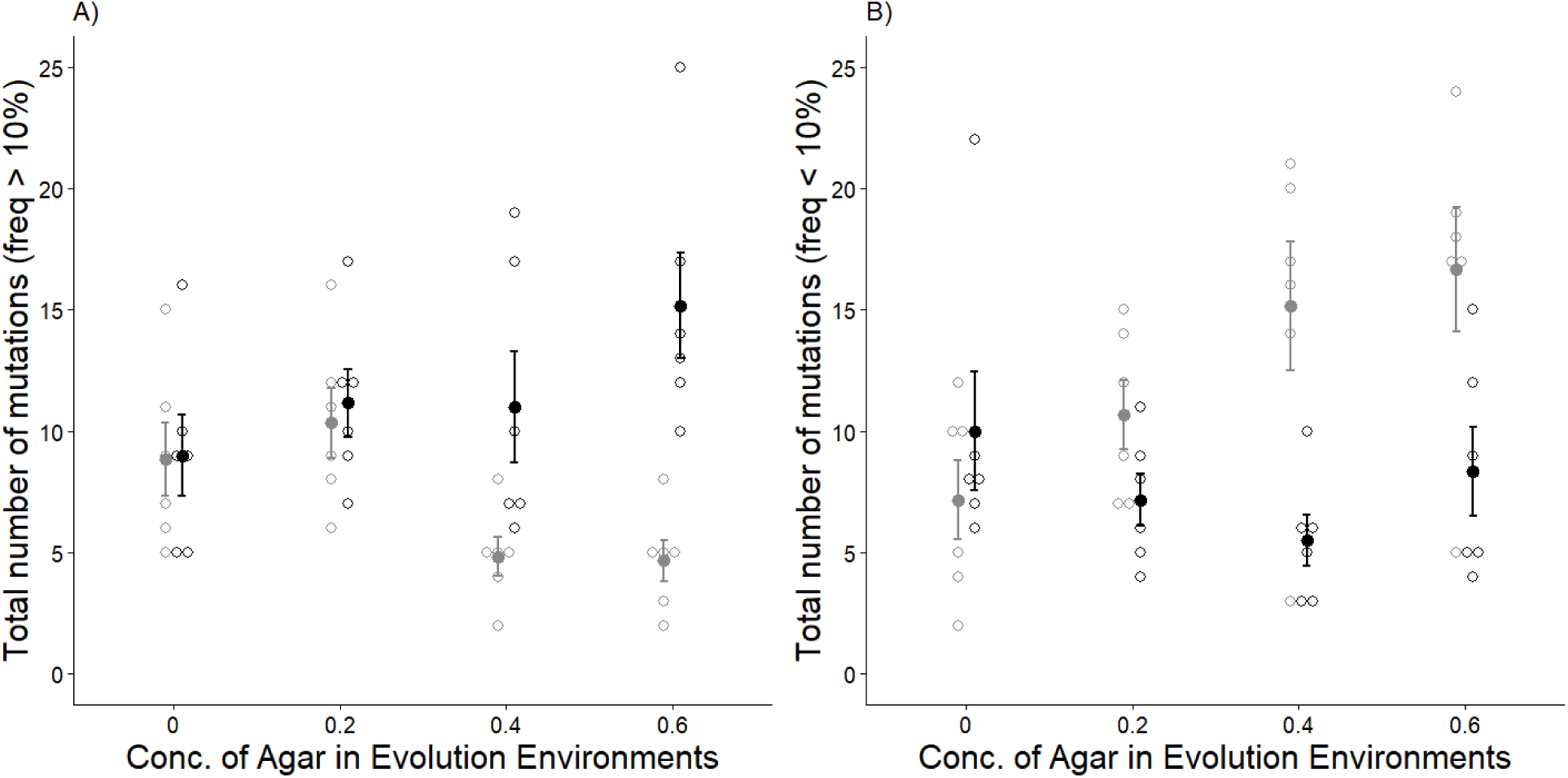
Total number of A) high frequency mutations (present in > 10 % of the population), and B) low frequency mutations (present in <10 % of the population), evolved in the absence (grey) and presence (black) or antibiotic, and across different concentrations of agar. Scattered blank circles represent the change of resistance of each replicate, filled circles with error bars represent the mean of six replicate populations +/- 1 SE).

The number of *low frequency* mutations was also significantly affected by agar concentration, antibiotic and their interaction (Fig. 3B; permutation ANOVA, antibiotic x agar concentration, P = 0.002, see Table S3). However, these mutations increased with increasing agar concentration in antibiotic-absent populations, and did not change significantly across agar concentration in antibiotic-present populations. In medium (0.4 %) and high agar concentration (0.6 % agar) environments, the number of low frequency mutations in antibiotic-absent evolved populations is higher than the antibiotic-present evolved population, the opposite of the high frequency mutations. We observed similar patterns and significant differences when we focus the comparison on non-synonymous mutations alone (Fig. S6, see Table S3)

We quantified the degree of parallel evolution at the gene level using the Jaccard Index (J) and compared J values for pairs of populations within each evolution environment, i.e. antibiotic present/ absent and agar concentration. Across all environments, observed J values (J_obs_) far exceed the null expectations (J_null_) assuming mutations arising randomly across the genome at the same rates as in our observed data (mean J_null_ = 0.0016 +/- 7.7×10^-5^ SE; mean J_obs_ = 0.0580 +/- 0.0048). There were also significant differences between J-values across treatments. We saw that for populations evolved in the presence of antibiotics, the degree of parallel evolution significantly increased with agar concentration, while for populations evolved with antibiotics, the degree of parallel evolution significantly decreased with agar concentration (Fig. 4; ANOVA, agar conc. x antibiotics P <0.001, see Table S4). This pattern remains consistent, even when we restrict our measures of parallel evolution to only high frequency mutations (>10% frequency) or only low frequency mutations (<10% frequency), as well as focusing on just the nonsynonymous mutations (Fig. S7; Table S4).

**Figure 4:**
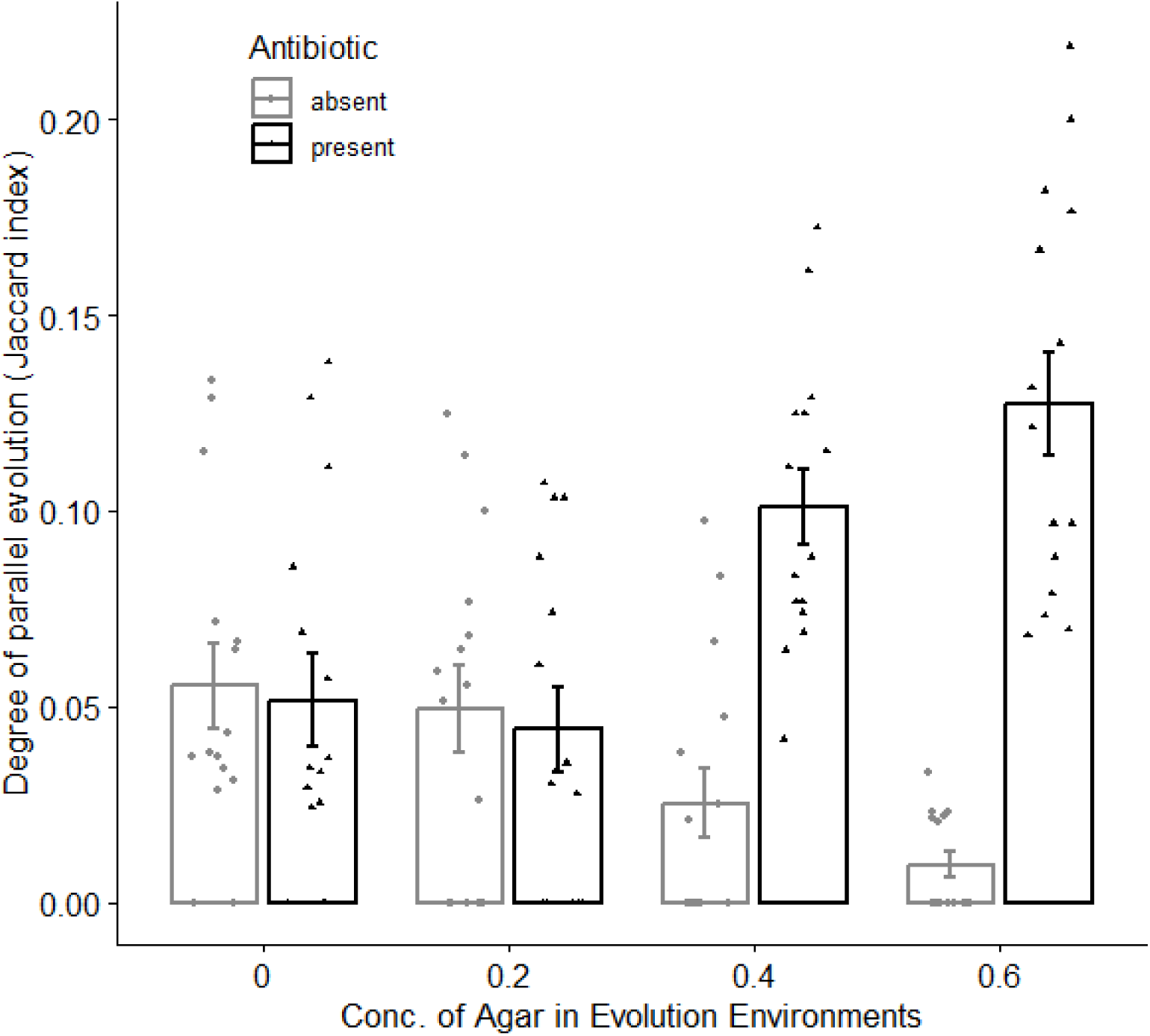
Degree of parallel evolution or pairwise gene-level similarity, quantified by the Jaccard index for evolved populations across different agar conc. environments in the absence (grey) and presence (black) of antibiotic. Bars and error bars indicate mean +/- standard error and point indicate each pairwise J value for that environment.

## Discussion

We tested the effects of agar concentration on the evolution of populations of *Pseudomonas aeruginosa* (PA) growing in the presence versus absence of a sub lethal concentration of the antibiotic, ciprofloxacin. The presence of antibiotics drove the rapid evolution of increased resistance to the antibiotic; however, the agar concentration and so degree of spatial structure in the environment did not significantly affect the level of evolved resistance. The degree of adaptation (i.e. relative fitness in the evolution environment) of the evolved populations, however, did vary with agar concentration, decreasing as the agar concentration increased and the presence of antibiotics increased the strength of this effect. We also observed impacts of agar concentration and antibiotics on evolved differences in population genomics. In the presence of antibiotics, the number of evolved high-frequency (putatively beneficial) mutations increased significantly with increasing agar concentration. In contrast, in the absence of antibiotics the number of high frequency (putatively beneficial) mutations decreased, while the number of low frequency mutations increased, with increasing agar concentration.

### Spatial structure slows the rate of adaptation

As agar concentration and so the degree of spatial structure increased in these populations, the rate of evolutionary adaptation (i.e. relative fitness, measured as change in relative growth rate in the evolution environment) decreased in both the presence and absence of antibiotics (Fig. 1). This observation fits with theoretical expectations for evolution in rugged fitness landscapes (e.g. Gordo and Campos 2006), and extends observations from previous experiments comparing populations grown in the extremes of very high structure (e.g. solid agar) versus no structure (e.g. well-mixed liquid) (France and Forney 2019), as well as multi-patch metapopulations with experimenter-mediated movement (Kryazhimskiy et al. 2012). Our study shows the first experimental evidence for a gradual shift in rate of adaptation across a range of intermediate degrees of spatial structures in a continuous-space environment. Another prediction arising from theoretical models is that while adaptation is expected proceed more slowly in spatially structured environments, eventually the extent of adaptation in structured environments is expected to exceed that of populations evolving in unstructured environments (Gordo and Campos 2006; Martens et al. 2011). We see no evidence for this pattern in our experiment, but we suggest that the populations reported on here may have simply not evolved for long enough for these longer-term patterns to emerge.

We note that in our experiment, all replicate populations underwent a daily mixing and population bottleneck event as part of the serial transfer regime. Despite this daily mixing, it is clear that the variation in structure experienced between the daily transfer events was still enough to drive important differences in evolutionary outcomes. (Habets et al. 2006) observed that the rate of adaptation in *E. coli* populations was slower in unmixed structured environments compared to structured environments with periodic mixing. Thus, the clear effects of agar concentration, and so spatial structure, observed in this study are likely to be further amplified if one were to use an experimental design that eliminated daily mixing.

Populations in all treatments of our experiment had significantly increased fitness by the end of the experiment; however, populations evolved in the absence of antibiotics had significantly lower fitness improvements compared to in the presence of antibiotics. This is expected, as populations evolving in the presence of antibiotics were under much stronger selection - antibiotic exposure reduced their initial growth rates by approximately 50%, by design. In the antibiotic-present populations, differences in agar concentration also had a larger impact on the rate of adaptation, suggesting that spatial structure may have particularly important impacts on evolutionary dynamics when selection is strong.

### Evolution of antibiotic resistance

Agar concentration affected many aspects of evolutionary dynamics in this study, but we detected no significant effect of agar concentration on antibiotic resistance, quantified by population-level MIC. It is possible that the absence of significance is simply due to low statistical power arising from the high variation in evolved MIC measured across the six replicate populations per treatment – certainly the trend suggests that evolved MIC decreases with increasing agar concentration, similar to observed significant effect of agar concentration on fitness. However, the absence of a significant effect may also arise because we held antibiotic selection pressure constant across environments (by design), selecting antibiotic concentration for each agar concentration such that it produced an approximately 50 % reduction in ancestral growth rate.

Despite no significant differences in the level of evolved resistance at the population level, we observed much variation in the underlying genetic routes to antibiotic resistance across the evolved populations. We observed mutations in six different previously identified antimicrobial resistance genes (mexR, mexS, mexZ (also referred to as amrR), nalC, nalD, and nfxB). Previous studies suggest two key ways that PA evolves resistance against ciprofloxacin and those are: 1) specialized quinolone resistance mutations, and 2) upregulation of efflux pumps (Hancock and Speert 2000; Breidenstein et al. 2011). In our study, we see mutations involved in upregulation of efflux pumps. Along with these key mechanisms, many other genes have pleiotropic effects on resistance depending on environment, and so the impacts of those diverse mutations on resistance are typically unclear (Breidenstein et al. 2008). Even within the same population, a diversity of clones with unique combinations of interacting mutations may result in wide variation in resistance (Rehman et al. 2019). In this study we measured resistance of populations (as opposed to single clones), and so we report population-level outcomes of a diversity of potentially interacting antibiotic resistance mechanisms present in that population. To give an example of the resistance diversity that arose: in one population (evolved in 0% agar), we observed mutations in three different known AMR genes: a repressor of the mexXY multidrug resistance efflux system (mexZ), and two different repressors of the mexAB multidrug resistance efflux system (mexR and nalC). We note also that there was no evidence of an evolved cost of resistance in our populations – growth rate measured in the absence of antibiotic did not correlate (negatively or otherwise) with evolved resistance (Fig. S4) (Lenormand et al. 2018). Future work will explore how the genetic diversity observed both within and between populations may be linked to variation in antibiotic resistance as well as other phenotypes, across different spatial structures.

### Adaptation to xylose

Many Pseudomonads (PA included) are reported to grow very slowly, or even below detection limits, when xylose is their sole carbon source (e.g. Liu 1952)). In this study, we chose to use a xylose-limited minimal media for our experimental populations, aiming to exert significant selection pressure on our experimental populations (even in the absence of antibiotic), and so allowing us to observe adaptive evolution over a relatively short period of time (as in Bailey and Kassen 2012). This experimental design choice proved successful, as we did observe a significant increase in fitness in populations across all environments. We expect that if a more-preferred carbon source had been used, we may have observed much less or even no adaptation (as well as parallel evolution) in the antibiotic-absent populations.

While evolutionary adaptations in our experimental populations increased fitness in the presence of xylose, we are unable to identify the specific mutations that impacted xylose metabolism. The PA14 genome only contains a single gene that codes for a protein known to be involved in xylose metabolism, and that is xylB, predicted to code for xylulokinase. We did not observe mutations in this gene in any of the evolved populations in this experiment. Thus, while we are unable to confirm which, if any, genes contributed to adaptation to xylose, we suggest some of the best candidates are ones predicted to code for proteins involved in transport that come up in multiple times in both antibiotic present and absent populations. For example, PA14_45060 – a predicted urea transport gene with observed mutations in seven antibiotic-absent populations and twelve antibiotic-present populations, as well as PA14_22650 – an ABC-type transport auxiliary lipoprotein family protein in which we observe fixed mutations in two antibiotic-absent populations and two antibiotic-present populations. We might also expect genes involved in motility to have general impacts on fitness across both antibiotic-present and -absent populations; however, we did not observe mutations in known motility-related genes arising across multiple evolved populations in this experiment.

### Population genomic changes

In this study, we observed that genomic patterns differed in populations evolved in different concentrations of antibiotics and these patterns also depended on the presence/ absence of antibiotics. As discussed previously, populations evolved in the presence of antibiotics were subject to stronger selection, resulting in faster rates of adaptation. Consistent with observed larger fitness increases, we saw a significantly larger number of putatively beneficial mutations in the evolved antibiotic-present populations. By “putatively beneficial”, we mean those mutations that appear to have escaped drift and are rising/ have risen to a relatively high frequency in the population. We identify these as mutations that are present at > 10 % in a population (Fig. 3A), and those mutations that are both high frequency and non-synonymous (see Fig. S6A). In other words, stronger selection imposed by the presence of antibiotics resulted in a larger number of putatively beneficial mutations escaping drift, and so driving larger total fitness gains in those populations.

Since increased agar concentration decreased the rate of adaptation in our experiment, and so we might expect to see evidence of this at the genomic level via a decrease in the number of putatively beneficial mutations. However, in this case what we see is a bit more complicated. For populations evolved in the absence of antibiotics, we do observe a decrease in the number of putatively beneficial mutations with increasing agar concentration as expected (Fig. 3A). However, in the presence of antibiotics, the number of putatively beneficial mutations actually *increased* with increasing agar concentration (Fig. 3A). We suggest that this increase arose from increased diversity of putatively beneficial mutations in spatially structured environments. Clonal competition likely increases with spatial structure, and so while beneficial mutants may arise and escape drift relatively quickly due to strong selection for antibiotic resistance, multiple beneficial mutants are then maintained over an extend period as they compete with each other for fixation. Previous experimental studies have also observed a significant increase in diversity in structured evolving populations (e.g. (Habets et al. 2006). It is also possible that the multiple high frequency mutants coexisting in the antibiotic-present spatially-structured environments represent stable diversity maintained through frequency dependent selection (e.g. as observed in (Rainey and Travisano 1998). A detailed characterization of the fitness and genotype of individual clones, beyond the scope of this study, would be required to distinguish these possibilities.

If we focus on the low frequency mutations (< 10 % of the population, but above our detection limit of 5 %), in general, we see fewer mutations in the antibiotic-present populations (Fig. 3B). We suggest that this pattern arises from a combination of 1) strong negative selection driving the relatively rapid loss of any significantly deleterious mutations that arise, and also 2) strong positive selection driving relative rapid increases in the frequency of beneficial mutations, shifting to frequencies greater than 10 %. In summary, we expect strong selection for antibiotic resistance to reduce the number of low frequency mutations in general. On the other hand, weaker selection in the absence of antibiotics likely allows neutral and nearly neutral mutations to be maintained in the populations for longer, and even more so in populations where the environment is spatially structured.

### Parallel evolution

In the presence of antibiotics, we saw that the degree of parallel evolution (measured at the gene-level) was significantly higher overall and increased with increasing agar concentration (Fig. 4). This was expected as antibiotics typically impose strong selection pressure leading to highly repeatable evolutionary outcomes (Santos-Lopez et al. 2021). The observed increase in parallel evolution with agar concentration in the antibiotic present environments is also expected as clonal competition increases with spatial structure. These results fit with previous observations from structured natural and experimental populations (e.g. biofilms: Steenackers et al. 2016, heterogeneous lab environments: Bailey et al. 2015; but Schick et al. 2022 a for counter example). However, in the absence of antibiotics the degree of parallel evolution, unexpectedly, decreased with agar concentration. In the antibiotic-absent high agar concentration populations, there was much less adaptive evolution overall and so we suggest that the decrease is simply a reflection of a much smaller number of adaptive mutations having arisen. Given more time for adaptive evolution in the antibiotic-absent populations, we suggest that the expected pattern of increased parallelism with increased agar concentration might start to emerge.

### Evolution of motility

Our aim in this study was to test the impact of agar concentration on evolution and so exploring the impact of differences in average movement rates (among other things) of bacteria through those environments and in turn their local interaction neighborhood. However, it is very possible that movements rates could also evolve in these populations, and so further shift the effective degree of spatial structure. Evolutionary changes in motility have been observed in a number of bacterial experimental evolution studies conducted in well-mixed liquid media, often via loss-of-function mutations in flagella-related genes (e.g. Barrick and Lenski 2009; Bailey et al. 2015). To look for evidence of this potential shift, we tested motility rates of our evolved populations in a common environment of 0.4 % agar (see Fig. S8). We saw that motility rate significantly decreased in populations evolved in the presence of antibiotics (Table S5) and while it did tend to also be lower in the antibiotic-absent populations, this was not a significant effect. We suggest that this may be because in the absence of the antibiotic (a less challenging environment), motility (which can be energetically costly for bacteria, Zhu and Gao 2020) has less of an impact on fitness and so selection is less efficient at reducing that cost.

Differences in motility have been reported to affect the ability of bacteria to respond to environmental stressors, such as antibiotics (Sun et al. 2018). PA in particular has been observed to shift its mode and rate of movement in response to structure in the environment, in particular viscosity (Sun et al. 2018), and both the mode and rate of motility can also evolve (Taylor et al. 2015). Adaptive evolution of motility *modes* to specific concentrations of agar may also contribute to the patterns of dispersal evolution we observe in the presence of antibiotics. Preliminary observations of colony dispersal shapes in the evolved populations from this study suggest that some populations may have evolved an increased tendency to swarm versus swim, compared to their ancestor. However, further detailed testing would be needed to confirm these patterns. It is also important to note that due to logistical constraints we conducted all our motility assays in 0.4 % agar, and so it is possible that evolved motility rate and mode changes are agar-concentration specific. Tests exploring evolved rate and mode of motility across a range of viscosities would allow for identification of just how specialized any motility adaptations are to a particular concentration of agar.

### No evidence of biofilm formation

In our study we focus on spatial structure generated by agar concentration in the environment, however biofilm formation in bacterial populations also generates spatial structure and can have important impacts on evolution (Santos-Lopez et al. 2019). In our experimental populations, however, we saw no evidence of biofilm formation. Biofilm formation in PA largely depends on quorum sensing, a small molecule regulated system that requires a high population density to initiate (Lee and Zhang 2015). In our study, we used a minimal media with xylose, an unpreferred nutrient source for PA, as the sole carbon source, and so the resulting population density was too low to expect successful quorum sensing and biofilm maintenance to have occurred.

### Conclusions

Spatial structure, a ubiquitous feature of the natural world, can have important impacts on evolution, from fitness to phenotype to population genomics, particularly within the context of antibiotic resistance evolution. The strong selection, arising from the presence of antibiotics in our experiment, is an important driver of these differences across different agar concentrations and so we suggest the general patterns described here may be applicable to microbial populations evolving in structured environments with other challenging stressors (e.g. high salt, pH). However, it is also likely that specific pleiotropic effects of motility-related genes on antibiotic resistance also play a role. Some genes have been shown to impact quinolone resistance as well as bacterial motility (e.g. *orfN;* Clarke et al. 2020; Laborda et al. 2022) and shifts in motility modes and rates are known to be affected by viscosity and nutritional signals in the environment (Harshey 1994). To understand the degree to which these evolved patterns are antibiotic resistance specific, future work will be aimed at exploring the phenotypic and genomic links between evolved motility and antibiotic resistance, at both the population- and clone-level, across a range of structured environments.

## Data availability

Sequence reads will be deposited in NCBI. Phenotypic and summary data along with R-scripts are available on request from the authors.

