## Supplementary figures and tables for "Agar concentration affects the evolution of antibiotic resistance and population genomics in experimental populations *Pseudomonas aeruginosa*"

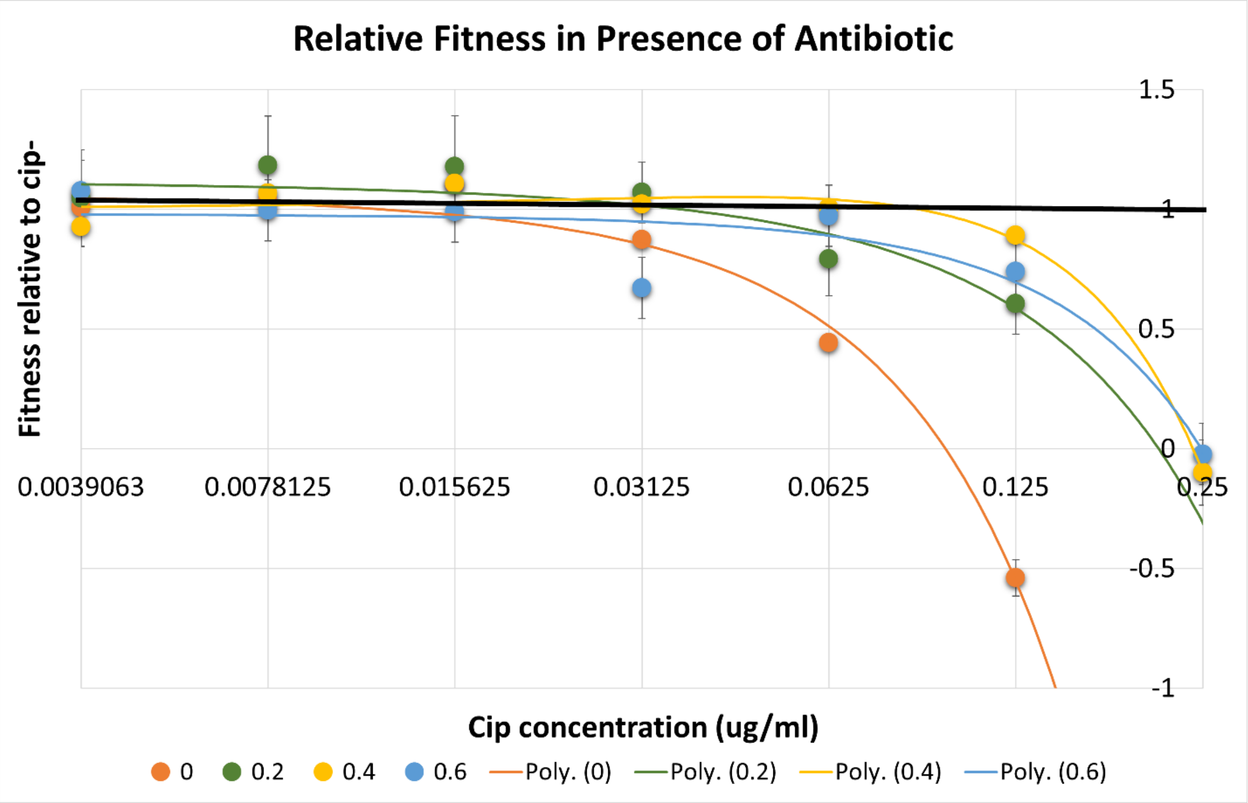


**Figure S1:** Relative fitness of PA ancestor in different concentrations of ciprofloxacin in M9 salts + xylose media with 0 %, 0.2 %, 0.4 %, and 0.6 % agar. X axis represents the ciprofloxacin concentrations; Y axis represents relative fitness measured as growth rate of bacteria in the presence of ciprofloxacin divided by the growth rate of the bacteria in the same media without ciprofloxacin.

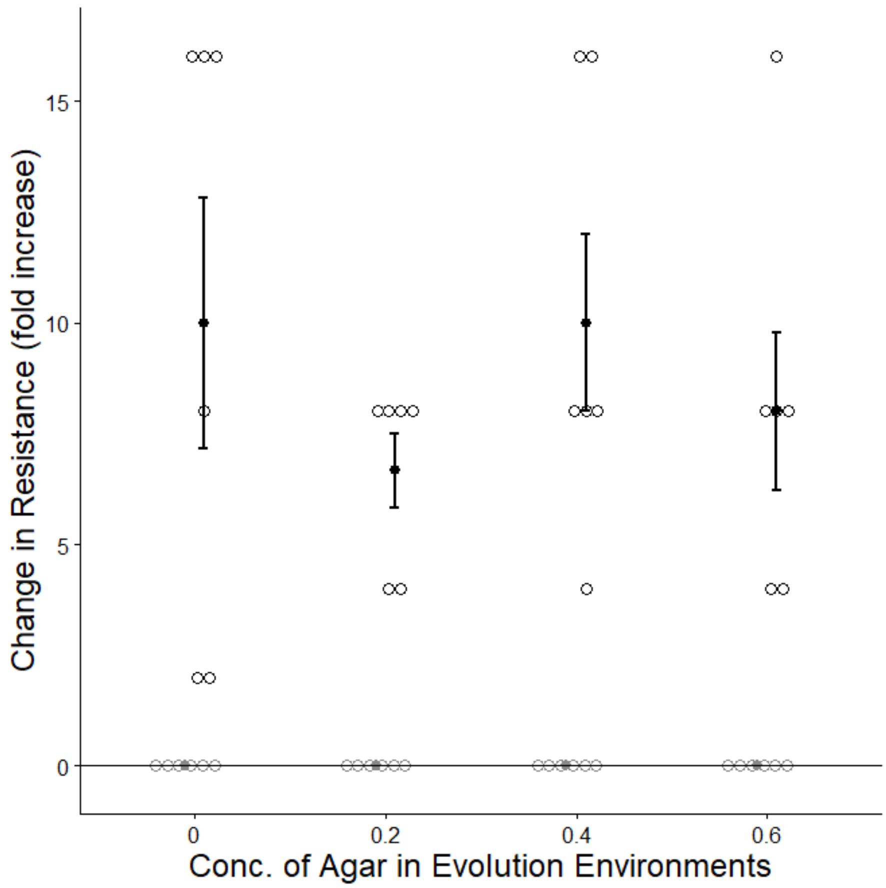


**Figure S2:** Change in resistance to ciprofloxacin of evolved populations relative to ancestor in Muller Hinton broth (MHB). Scattered blank circles represent the change of resistance of each replicate, dark points with error bars represent the mean of six replicate populations +/- 1 SE.

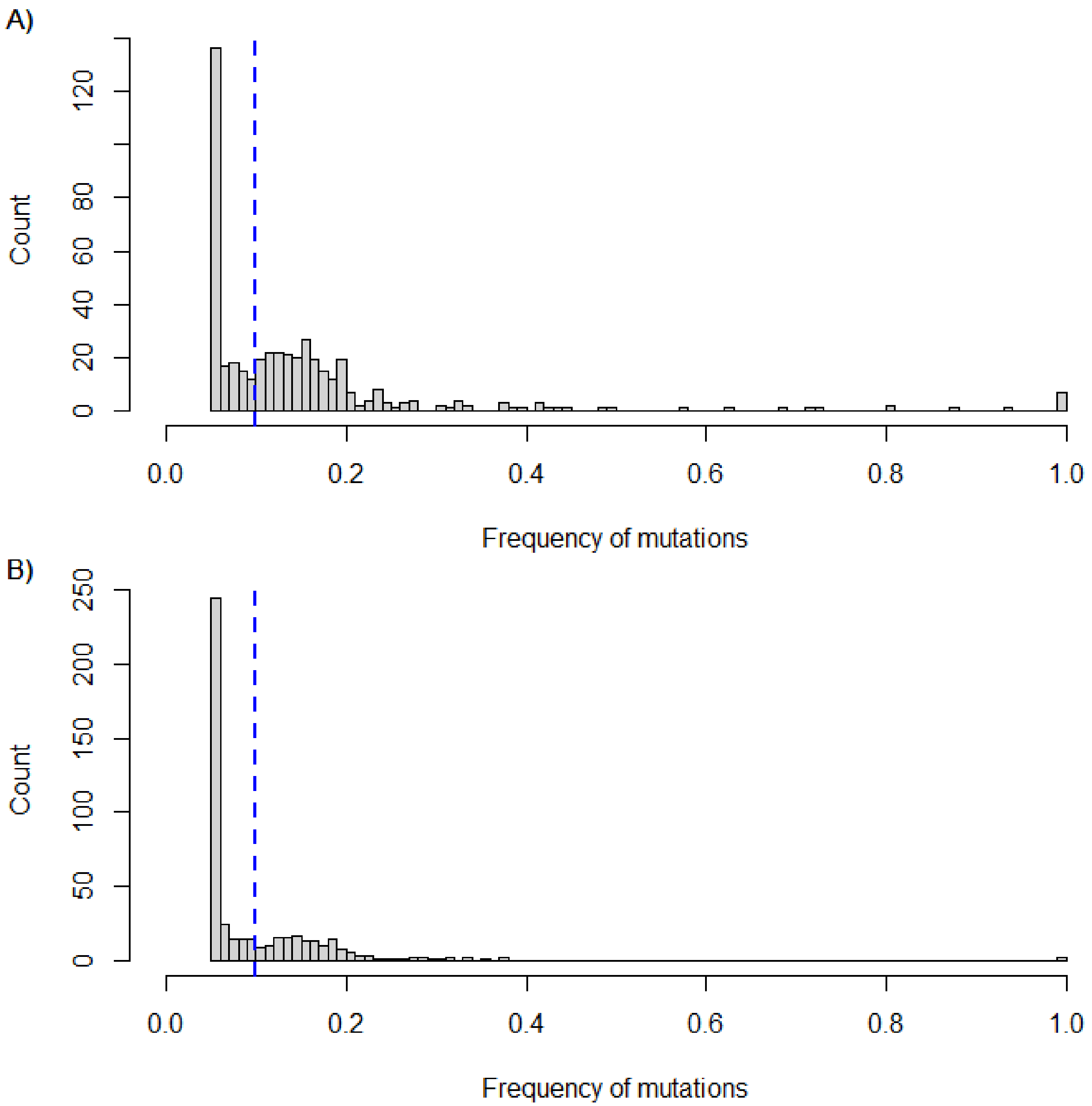


**Figure S3:** Distribution of mutations frequencies in all populations evolved in A) the presence of antibiotics, and B) the absence of antibiotics. The minimum frequency detected (due to limitations of the methods) is 0.05. The vertical blue dashed line shows the cut-off of 10% that we chose to group mutations into “high” and “low” frequency.

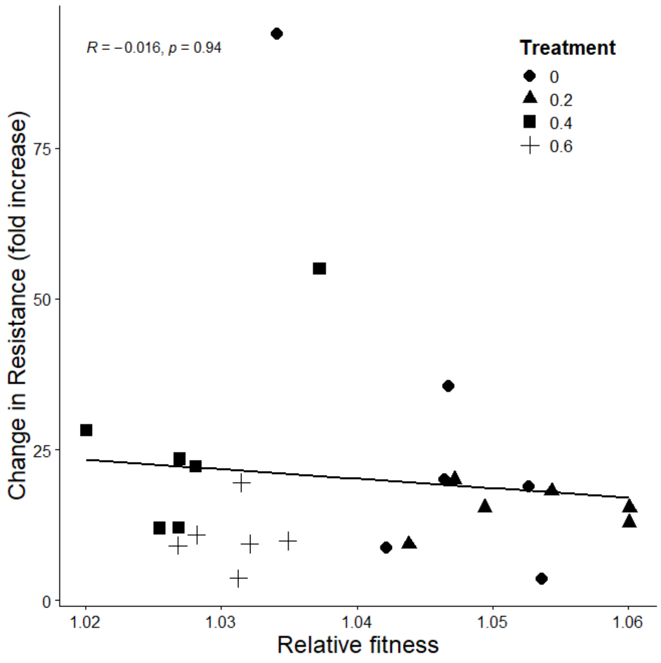


**Figure S4:** Increase in resistance to ciprofloxacin relative to the ancestor over relative fitness for each evolved population tested in the spatial structure they evolved in but without ciprofloxacin. Data presented here is only for the antibiotic-present evolved populations. Line shows the best fit regression, slope of this line is not significantly different from 0; linear regression P>0.05).

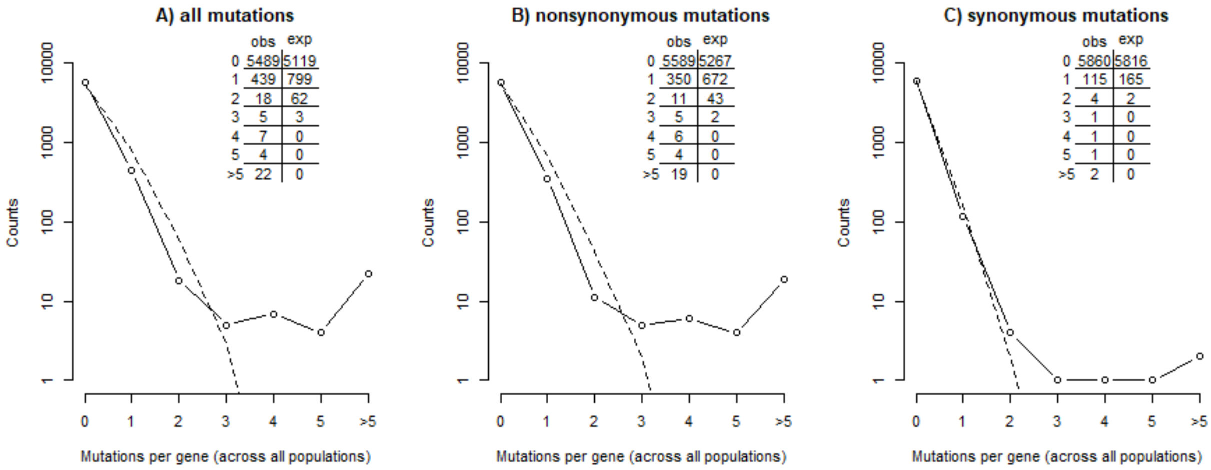


**Figure S5:** The distribution of number of hits per gene pooled across all 48 evolved populations, including A) all types of mutations, B) nonsynonymous mutations only, and C) synonymous mutations only. Circles joined by solid lines represent observed data and dashed lines represent the null expectation assuming mutations arise via a Poisson random process with mutation rate equal to that of the observed data. Inset tables show the same data as on plot, displayed numerically. Kolmogorov-Smirnov tests comparing observed distributions to the expected null distribution (assuming Poisson) all indicate that there is a significant difference between the observed and expected (P < 0.0001 for all three comparisons)

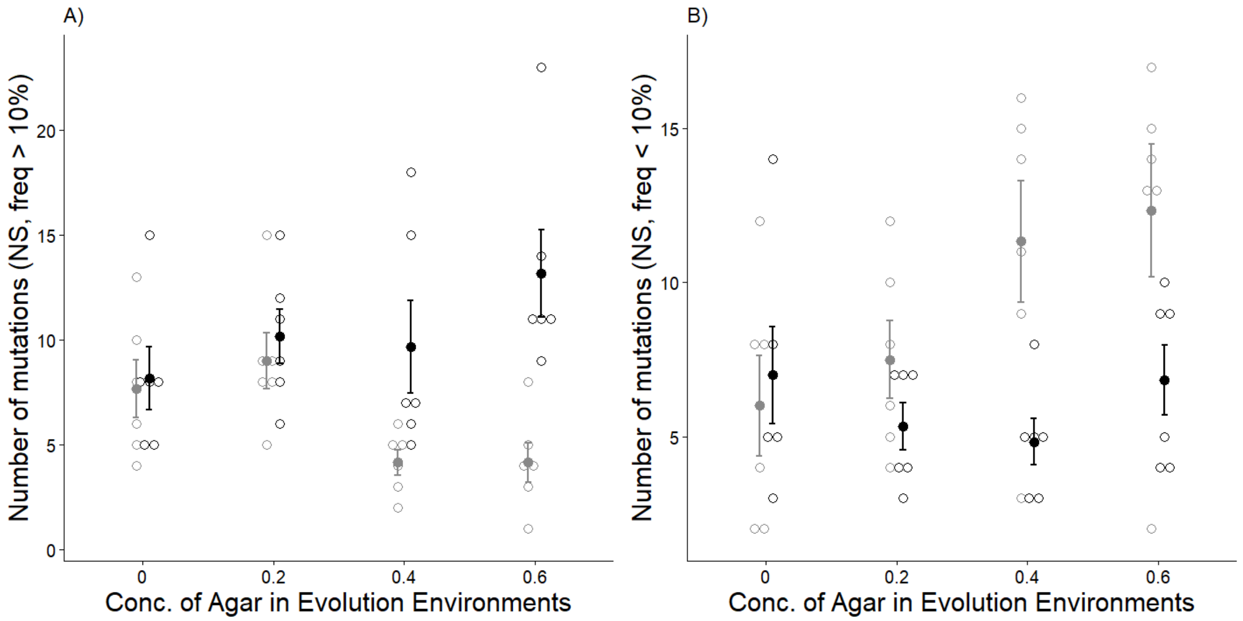


**Figure S6:** Total number of A) high frequency nonsynonymous mutations (present in > 10 % of the population), and B) low frequency nonsynonymous mutations (present in <10 % of the population), evolved in the absence (grey) and presence (black) or antibiotic, and across different concentrations of agar. Scattered blank circles represent the change of resistance of each replicate, filled circles with error bars represent the mean of six replicate populations +/- 1 SE).

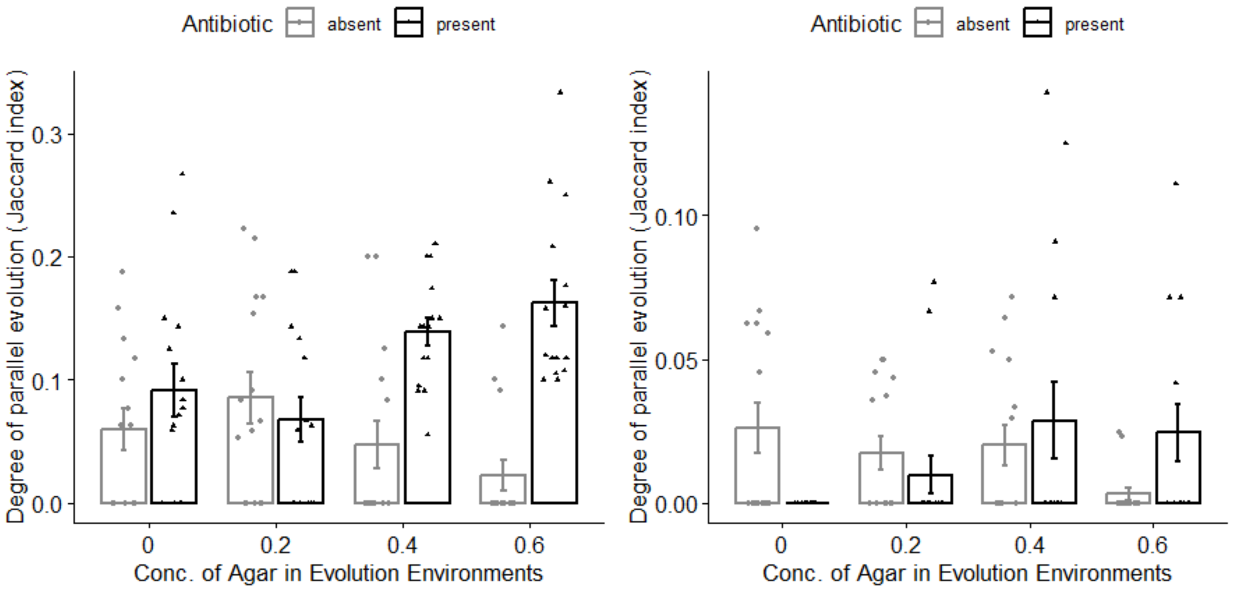


**Figure S7**: Degree of parallel evolution or pairwise gene-level similar, quantified by the Jaccard index for evolved populations across different concentrations of agar in the presence (black) and absence (grey) of antibiotic. Left panel shows the Jaccard index calculated with only mutations with greater than 0.1 frequency in a population; right panel shows the Jaccard index calculated with only mutations with less than or equal to 0.1 frequency in a population.

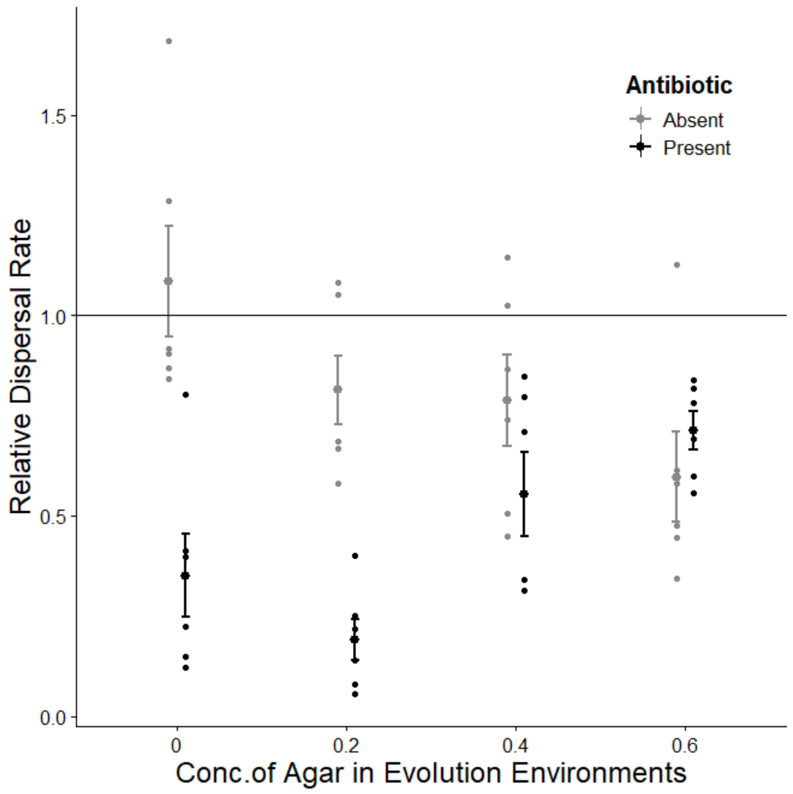


**Figure S8:** Relative dispersal rate of populations evolved with antibiotic absent (gray) and present (black) across different concentrations of agar, all assayed in semi-solid (0.4 %) King B media. Scattered points represent six replicate populations, while points with error bars represent the mean +/- 1 SE for each treatment. The solid horizontal line at y = 1 indicates the dispersal rate of the ancestor.

Here, motility rates were estimated in semi-solid KB media with 0.4% agar by inoculating Petri dishes with evolved populations and an ancestral strain. Each dish was pricked with a needle, then incubated at 37°C. After 24 hours, colony areas were photographed and measured using ImageJ software. To distinguish motility from growth, population sizes were estimated by measuring optical density at 0 and 24 hours. The dispersal rate relative to the ancestor (relDev) was calculated using:

$$relD_{ev} = ( A_{ev} / \Delta OD_{ev} ) / ( A_{anc} / \Delta OD_{anc} )$$

where *A* is the colony area, ($\Delta$OD) is the estimated change in population density, and subscripts *ev* and *anc* refers to the evolved population and ancestor, respectively.

Dispersal rates decreased in evolved populations compared to the ancestor (fig. S5). Antibiotics significantly reduced dispersal rates (permutation ANOVA, N = 1000, $P < 0.001$), and this effect varied with spatial structure (interaction antibiotic x structure, $P < 0.001$). Populations evolved with antibiotics had significantly lower dispersal rates in three out of four structures, while those without antibiotics did not show a significant decrease (see table S5).

### Supplementary Tables

**Table S1:** Results of ANOVA testing the effects of agar concentration (0, 0.2, 0.4, 0.6 %), antibiotic (present/ absent), and their interaction on relative fitness.

| **Factor** | P-value |
|---|---|
| **Agar conc.** | 0.009 *** |
| **Antibiotic** | <0.0001 *** |
| **Agar conc. x Antibiotic** | 0.006 *** |

**Table S2:** Mean number of mutations per population, grouped by treatment and mutation type. Here, nonsynonymous counts include all genic mutations that are not synonymous (i.e. indels, nonsense).

| | | All mutations (mean per population) | | | | | |
|---|---|---|---|---|---|---|---|
| | | nonsynonymous | | synonymous | | intergenic | |
| | -cip | mean | SE | mean | SE | mean | SE |
| agar conc. | 0 | 13.7 | 1.1 | 2.3 | 0.5 | 12.5 | 1.3 |
| | 0.2 | 16.5 | 1.7 | 4.5 | 0.8 | 11.0 | 1.4 |
| | 0.4 | 15.5 | 2.0 | 4.5 | 0.7 | 7.0 | 0.5 |
| | 0.6 | 16.5 | 2.5 | 4.8 | 0.9 | 6.2 | 2.1 |
| | +cip | mean | SE | mean | SE | mean | SE |
| agar conc. | 0 | 15.2 | 1.5 | 3.8 | 1.0 | 7.7 | 1.2 |
| | 0.2 | 15.5 | 1.0 | 2.8 | 0.5 | 11.5 | 1.1 |
| | 0.4 | 14.5 | 1.8 | 2.0 | 0.3 | 12.0 | 1.7 |
| | 0.6 | 20.0 | 1.8 | 3.5 | 0.7 | 14.2 | 2.2 |
| | All | 15.9 | 0.6 | 3.5 | 0.3 | 10.3 | 0.6 |
| | | Fixed mutations (mean per population) | | | | | |
| | | nonsynonymous | | synonymous | | intergenic | |
| | -cip | mean | SE | mean | SE | mean | SE |
| agar conc. | 0 | 0.2 | 0.4 | 0.0 | - | 0.3 | 0.5 |
| | 0.2 | 0.2 | 0.4 | 0.0 | - | 0.2 | 0.4 |
| | 0.4 | 0.0 | - | 0.0 | - | 0.8 | 0.7 |
| | 0.6 | 0.0 | - | 0.0 | - | 0.0 | - |
| | +cip | mean | SE | mean | SE | mean | SE |
| agar conc. | 0 | 0.2 | 0.4 | 0.0 | - | 0.0 | - |
| | 0.2 | 0.7 | 0.5 | 0.0 | - | 0.3 | 0.5 |
| | 0.4 | 0.5 | 0.5 | 0.0 | - | 0.7 | 0.5 |
| | 0.6 | 0.0 | - | 0.0 | - | 0.0 | - |
| | All | 0.2 | 0.4 | 0.0 | - | 0.3 | 0.5 |

**Table S3:** Results of ANOVA testing the effects of agar concentration (0, 0.2, 0.4, 0.6 %), antibiotic (present/ absent), and their interaction on the total number of a) all mutations and b) just nonsynonymous mutations, in two different frequency classes – high and low.

| | **All Mutations** | | | **Nonsynonymous Mutations** | |
|---|---|---|---|---|---|
| | **High Frequency (>10%)** | | **Low Frequency (<10%) (Permutation ANOVA, N=1000** | **High Frequency (>10%) (Permutation ANOVA, N=1000** | **Low Frequency (<10%) (Permutation ANOVA, N=1000** |
| | **F- Value** | **P-Value** | **P-Value** | **P-Value** | **P-Value** |
| **Agar conc.** | 0.0011 | 0.9740 | 0.036 * | 0.931 | 0.022* |
| **Antibiotic** | 15.1214 | 0.0003 ** | 0.001 ** | <0.001*** | 0.004** |
| **Agar conc. x Antibiotic** | 12.7915 | 0.0009 ** | 0.002 *** | 0.004*** | 0.009** |

**Table S4: Results of permutation ANOVA testing the effects of antibiotics, agar concentration, and their interaction on parallel evolution measured by pairwise Jaccard index**. Jaccard index was calculated and compared for all mutations, nonsynonymous mutations (NS), synonymous mutations (S), high frequency mutations (>10%), and low frequency mutations (<10%). P-values <0.05 are indicated with *.

| | **P-values** | | | | |
|---|---|---|---|---|---|
| **Factor** | *All mutations* (N = 934) | *NS mutations* (N = 764) | *S mutations* (N = 170) | *High freq mutations* (N=423) | *Low freq mutations* (N = 511) |
| **Antibiotic** | <0.001* | <0.001* | 0.128 | <0.001* | 0.858 |
| **Agar conc.** | 0.172 | 0.135 | 0.016* | 0.636 | 0.380 |
| **Anti x Agar conc.** | <0.001* | <0.001* | <0.001* | <0.001* | 0.019* |

**Table S5:** Results of t-tests comparing evolved dispersal rate to that of the ancestor, across different degrees of agar concentration (0, 0.2, 0.4, 0.6 %) and antibiotic present vs. absent.

| **Relative dispersal rate of the evolved population** | | |
|---|---|---|
| **Agar conc.** | Antibiotic | |
| | Absent | Present |
| **0% agar** | 1 | 0.01187* |
| **0.2% agar** | 0.4936 | 0.00016*** |
| **0.4% agar** | 0.6436 | 0.22430 |
| **0.6% agar** | 0.1213 | 0.01535* |
